# Circadian clock genes *Bmal1* and *Per2* in the nucleus accumbens are negative regulators of alcohol-drinking behavior in mice

**DOI:** 10.1101/2023.02.24.529935

**Authors:** J. Herrera, M. Button, P. Doherty-Haigh, C. Goldfarb, N. Quteishat, S. Amir, K. Schoettner

## Abstract

Voluntary alcohol consumption is influenced by a variety of environmental and genetic factors, including circadian clock genes. Even though their sex-specific role in alcohol drinking was identified through selective ablation of *Bmal1* and *Per2* from neurons of the mouse striatum, the contribution of specific striatal subregions to the observed drinking behavior remains unclear. Thus, alcohol intake and preference was investigated in male and female mice with a conditional knockout of *Bmal1* and *Per2* from cells in the nucleus accumbens (Nac). Mood- and anxiety-related behaviors were assessed prior to alcohol drinking to exclude potential confounding effects of the animal’s behavioral state on alcohol consumption. Alcohol consumption and preference were increased in male and female mice with a conditional knockout of *Bmal1*, whereas the same effect was only found in males with a deletion of *Per2*. Because affective behaviors were only mildly influenced by the conditional gene knockouts, observed alcohol-drinking phenotypes can be directly associated with the Nac-specific clock gene deletion. The results thus suggest an inhibitory role of *Bmal1* and *Per2* in the Nac on alcohol consumption in male mice. In females, the inhibitory effect of *Bmal1* is strictly localized to the Nac, because striatal-wide deletion of *Bmal1* caused a suppression of alcohol consumption. This sex-dependent stimulatory effect of *Bmal1* on alcohol drinking is probably mediated through other striatal subregions such as the dorsal striatum.

## INTRODUCTION

Circadian clock genes are well recognized for their role in the generation and regulation of circadian rhythms in physiology and behavior as well as for their contribution to a broad range of pathophysiological conditions, including alcohol use disorder, when their function is compromised. The involvement of clock genes in the control of alcohol-drinking behavior is suggested by gene association studies in humans^1–3^ and supported by gene knockout studies in mice, which point to a causal link between loss of function in clock genes such as *Clock, Per1, Per2*, or *Rev-erb alpha* and changes in alcohol consumption^4–8^. Currently, however, brain regions and neural mechanisms at the intersection of these genes and alcohol consumption remain largely unexplored.

Selective deletion of *Bmal1* from medium spiny neurons (MSNs) of the striatum, a critical structure in the control of alcohol-drinking behavior, led to sexually dimorphic alterations in alcohol consumption – augmentation in males and suppression in females^9^. Like *Bmal1*, deletion of the clock gene *Per2* from MSNs led to heightened alcohol consumption in males. However, *Per2* deletion did not affect consumption in females. Together, these finding suggest that *Bmal1* and *Per2* in MSNs of the striatum normally restrain alcohol consumption in males, whereas, in females, a distinct *Bmal1*-dependent mechanism, promotes heightened alcohol intake^9^.

Although all major striatal subregions have been implicated in the control of alcohol consumption, the nucleus accumbens (NAc) is distinguished by its critical role in reward and appetitive motivation, and by its rich history of involvement in alcohol-drinking behavior^10,11^. Recent studies in mice have shown that suppression of the clock gene *Per1* or co-repression of the genes *Clock, Per1* and *Per2* in the Nac attenuates alcohol binge drinking^12^, whereas downregulation of *Clock* in the ventral tegmental area (VTA), a primary player in the mesolimbic dopamine reward circuit that projects to MSNs of the Nac, has been shown to augment alcohol intake in both male and female mice^8^. To further decipher the sexually dimorphic effects of striatal clock gene expression on alcohol consumption in mice, we used adeno-associated viral vectors to ablate *Bmal1* or *Per2* from cells of the Nac specifically. Alcohol intake and preference was investigated in male and female mice that have previously underwent behavioral assessment. Circadian clock genes in the Nac have been shown to play a role in the control of affective behaviors, which, in turn, could affect ethanol consumption^12–15^. We therefore evaluated anxiety- and depressive-like behavior in all mice prior to alcohol exposure, to rule out potential confounding effects of behavioral changes on ethanol consumption because of the Nac-specific clock gene manipulation.

## MATERIAL & METHODS

### Animals and housing conditions

Male and female C57BL/6J mice (12-16 weeks old) carrying floxed alleles of either *Bmal1 (Bmal1^fl/fl^*, Jackson Laboratory, stock number 007668) or *Per2 (Per2^fl/fl^*, European Mouse Mutant Archive, Strain ID: EM10599) were used in this study. Mice were housed individually in standard plastic cages in light- and soundproof boxes with room temperature and relative humidity varying around 21 ± 1°C and 65 ± 5%, respectively. Animals were kept under a 12-h/12-h light-dark schedule and had free access to food and water. The bedding of the cages was changed on a weekly basis. All procedures were reviewed and approved by the Canadian Council on Animal Care and the Concordia University Animal Care Committee (certificate number: 30000256).

### Mouse stereotactic surgery

Stereotactic surgeries were performed for Nac-specific delivery of adeno-associated viral vectors (AAV). Animals were anaesthetised with a mix of Ketamine and Xylazine and fixed to a stereotactic device. Mice with a conditional clock gene knockout (hereafter referred to as KO) received bilateral intra-Nac injections (anteroposterior: 1.6 mm posterior to bregma, mediolateral: 1.00 mm lateral to midline, dorsoventral: −4.7 mm ventral to the skull, 250 nl/ hemisphere) of AAVs expressing Cre-recombinase and Enhanced Green Fluorescence Protein (EGFP) (AAV2/5-CAG-CRE-EGFP, 1.0 x 10^12^ vg/ml, Canadian Neurophotonics Platform Viral Vector Core Facility, Quebec, Canada), whereas control animals (CTR) received AAVs expressing GFP only (AAV2/5-CAG-EGFP, 1.0 x 10^12^ vg/ml, Canadian Neurophotonics Platform Viral Vector Core Facility, Quebec, Canada). The spread and efficiency of AAV infection in the Nac were validated in all mice at the end of the study by immunofluorescence staining (Fig. 1).

**Figure 1:**
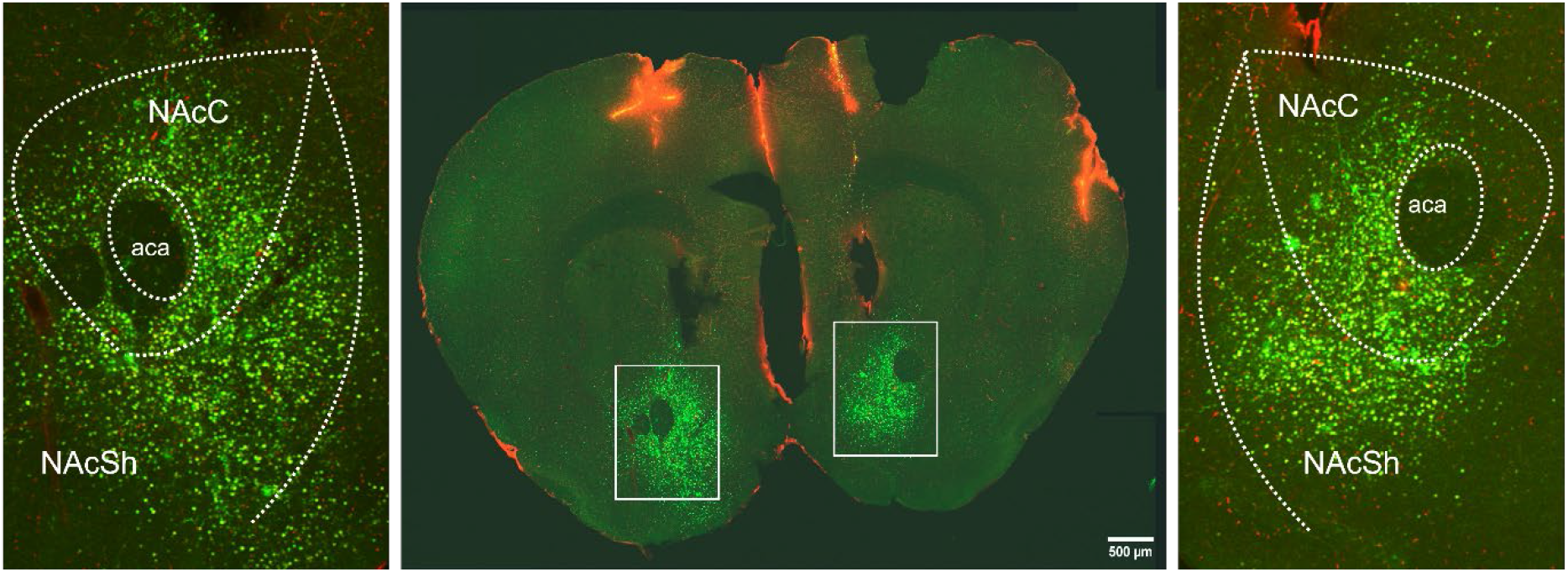
Representative image of GFP expression in the nucleus accumbens of mice receiving stereotaxic injections of AAV-CAG-Cre-EGFP viral vectors. GFP is strongly expressed in the nucleus accumbens core (NAcC) and shell (NAcS) region. Scale bar = 500 μm.

### Behavioral procedures, Ethanol consumption and sucrose preference

Three weeks after Nac-specific delivery of viral vectors, a sequence of behavioral tests was conducted to assess anxiety- and depressive-like behaviors. The order of the tests was fixed, starting with the elevated plus maze test (EPM) followed by the open field test (OFT), marble burying test (MBT), and tail suspension test (TST). Test were conducted during the light phase between Zeitgeber time (ZT) 2 and 6, except for the TST that was conducted at ZT8 (ZT0 refers to the time of light onset). The interval between experiments was at least 1 week. Experiments were carried out in a separate experimental room. Before each test, animals were habituated to the test room for 1 h. Details of the behavioral procedures and collected parameters are described elsewhere^16^. Voluntary alcohol consumption was studied subsequently using the intermittent two-bottle choice test as described previously^9^. In brief, mice were accustomed to drink water from two bottles for three days one week after the last behavioral test. They were then given access to one bottle of alcohol solution (15% v/v, in tap water) and one bottle containing tap water only every other day, for 11 sessions. The position of the alcohol and water bottles was altered each session to control for potential side preference. All mice were given access to two water bottles during the alcohol-off days. Body weights, alcohol intake (g/kg body weight/day), and alcohol preference (ml of alcohol solution consumed/total fluid intake) were measured and calculated at the end of each daily alcohol session.

A sucrose preference test was performed following Ethanol consumption measurements. The mice were first re-habituated to two bottles of tap water for three days, and then were given unlimited access to one bottle of tap water and one bottle of 0.25% sucrose solution in tap water for three consecutive days. The ratio of the sucrose solution consumed relative to the total fluid intake was used as a measure of taste preference.

### Immunofluorescence

Animals were deeply anesthetized by exposure to an atmosphere of isoflurane and perfused transcardially by cold saline (0.9% sodium chloride, pH 7.2) followed by paraformaldehyde solution (PFA, 4% in 0.1M phosphate buffer, pH 7.2) using an infusion pump. Brains were then dissected and postfixed in PFA for 22 – 24h at 4°C. 30μm coronal section of brain tissue were collected using a Leica vibratome and analyzed under a fluorescent microscope to validate region-specific viral vector delivery. Brain slices were stored at −20°C in Watson’s cryoprotectant (Watson et al., 1986) for immunofluorescence imaging. For this, free-floating sections previously kept in Watson’s cryoprotectant were rinsed for 10 minutes in phosphate buffered saline (PBS, pH 7.4), cover slipped with a mounting media containing DAPI and stored at 4°C until imaging under a Olympus 10FI confocal microscope. A ratio between GFP expressing cells over total number of DAPI stained cells was used to calculate rate of viral infection. Representative images were prepared using FIJI^17^ and Inkscape (Inkscape Project, 2020).

Subjects with missing, incomplete or off-region GFP signaling were excluded from the analyses of behavioral tests and ethanol consumption.

### Data analysis and statistics

The collected raw data from the behavioral tests and alcohol consumption experiment were further processed and analyzed using Prism 9 (GraphPad Software, San Diego, CA, USA). The outcomes were depicted as mean ± standard error of the mean (SEM). A two-way analysis of variance (ANOVA, viral vector, and sex effect) followed by Šídák’s multiple comparison test was conducted to analyze the behavioral tests. A two-way repeated-measures ANOVA (viral vector, and alcohol session effect) followed by Šídák’s multiple comparisons test was conducted to analyze alcohol consumption and preference in male and female mice with a Nac-specific clock gene manipulation. Averaged alcohol consumption and preference for knockout (KO) and CTR animals over 11 sessions were compared using an unpaired t-test. The level of significance was set at p ≤ 0.05.

## RESULTS

### Deletion of *Bmal1* and *Per2* in mouse Nac

Analysis of GFP and Cre recombinase expressing cells from all histologically validated surgeries revealed that 65-75% of Nac cells were infected with AAV expressing Cre and GFP (KO), or GFP only (CTR). This resulted in the following experimental and control groups: *Bmal1-KO* males (n=9), *Bmal1-CTR* males (n=7), *Bmal1-KO* females (n=8), *Bmal1*-CTR females (n=8), *Per2-KO* males (n=12), *Per2*-CTR males (n=13), *Per2*-KO females (n=9), *Per2*-CTR females (n=7).

### Effect of *Bmal1* deletion in the Nac on anxiety and depressive-like behaviors

Nac-specific deletion of *Bmal1* in male and female mice had no effect on behaviors displayed in the EPM and MBT (Fig. 2 A – C, G, Table 1). Although anxiety-related behavior displayed in the OFT wasn’t altered in general (Fig. 2 D + F, Table 1) because of the *Bmal1* ablation in the Nac, the knockout affected the latency to explore the center of the open field arena in male and female mice (Fig. 2 E, Table 1). Assessment of depressive-like behavior in the TST indicates a sex-specific effect of the deletion of *Bmal1* on the mobility time displayed by the animals. Whereas knockout males displayed significantly increased mobility in the test, the opposite trend was observed in females (Fig. 2 H, Table 1).

**Figure 2:**
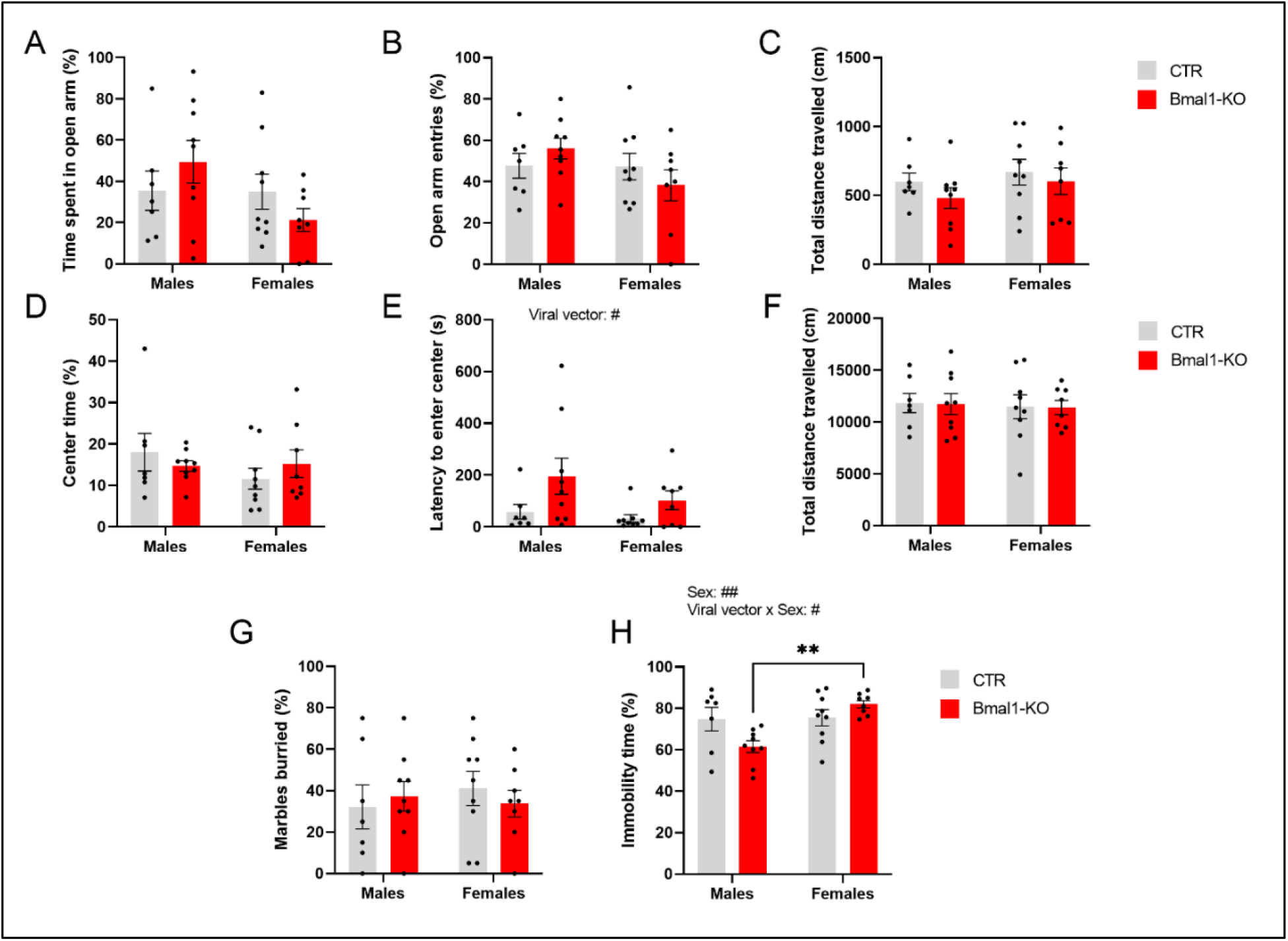
Anxiety- and depressive-like behavior in mice with a NAc-specific *Bmal1* knockout (n = 7 – 9 per experimental group and sex). The knockout had no effect on behaviors displayed in the elevated plus-maze (A – C) but increased the latency to approach the central area of the open field (E) whereas other parameter remained unchanged (D, F). No differences between groups were found in the marble burying test (G). Increased mobility in *Bmal1-KO* males was found when animals were suspended to bar in the tail suspension test (H). Results are depicted as mean ± standard error of the mean (S.E.M.). # … p ≤ 0.05, ## … p ≤ 0.05, two-way ANOVA. **… p ≤ 0.005, Šídák’s multiple comparisons test. Details of the statistical analysis are shown in Table 1.

**Table 1:**
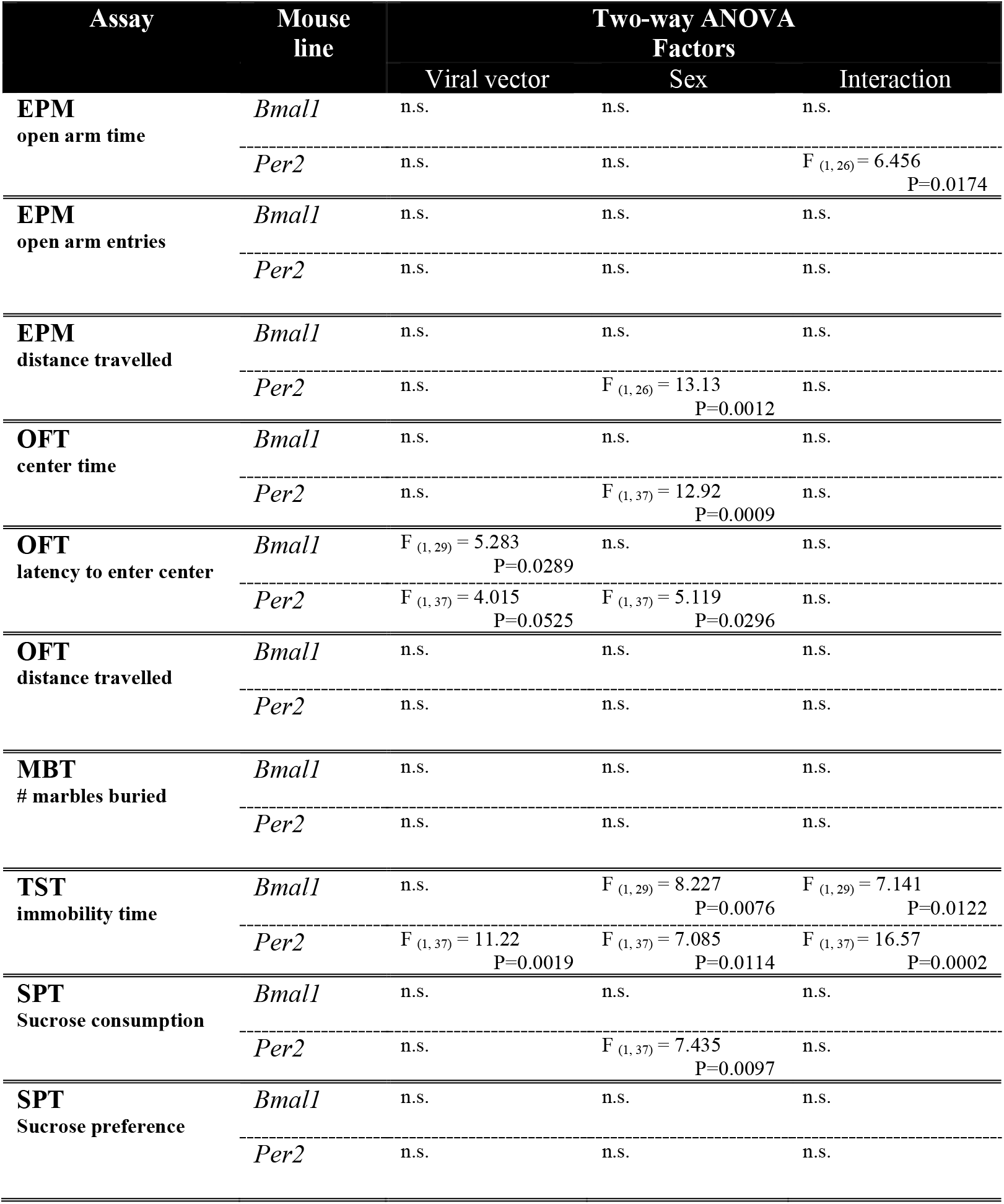
Results of behavioral testing.

### Effect of *Bmal1* deletion in the Nac on alcohol and sucrose consumption and preference

Deletion of *Bmal1* in the Nac significantly augmented daily alcohol intake and preference in both male and female mice although in females the increase in consumption was gradual (Fig. 3 A – D). Results from two-way repeated measure ANOVAs (Table 2) revealed a significant effect of the viral treatment on alcohol consumption and preference in both males and females as well as a significant effect of alcohol session on intake in males, and on intake and preference in females (Table 2). A significant viral vector x session interaction was noted in females but not in males (Table 2). On average, *Bmal1-KO* males consumed 4 g/kg more alcohol and showed 21% higher alcohol preference compared to CTRs (Fig. 3 A + C, right panel). Similarly, *Bmal1-KO* females drank significantly more than female CTR animals, 5 g/kg on average and preferred ethanol over water 18% more than CTR animals (Fig. 3 B + D, right panel). Body weights across the 11-sessions were not affected by deletion of *Bmal1* in either males or females (data not shown). To exclude the possibility that the observed changes in alcohol intake and preference seen in the *Bmal1-KO* mice were a result of a more general change in appetitive motivation, mice were given a sucrose preference test immediately following the last alcohol session. As shown in Fig. 3 E + F, in both males and females, deletion of *Bmal1* in the Nac had no effect on sucrose intake and preference (Table 1).

**Figure 3:**
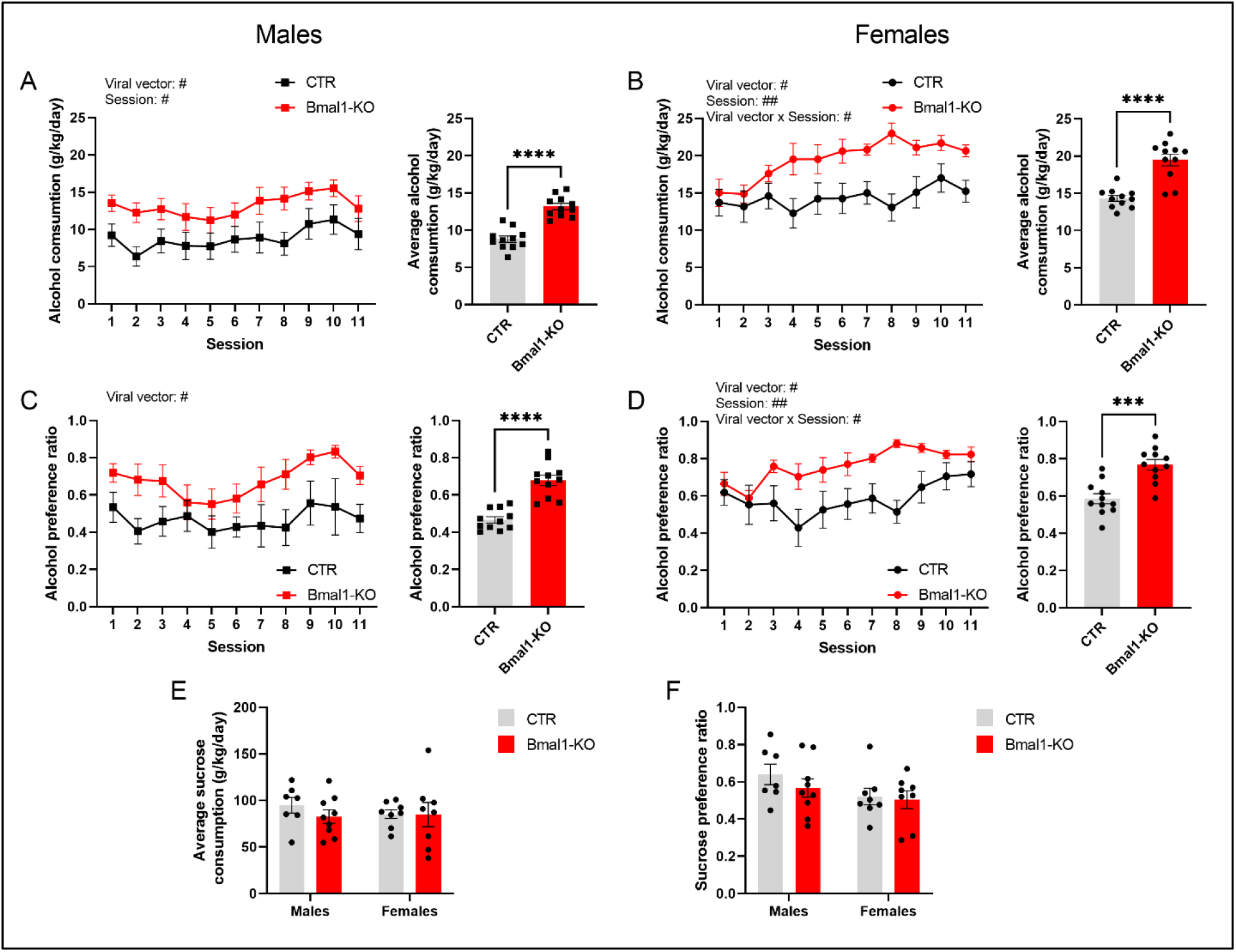
Alcohol and sucrose consumption and preference in mice with a *Bmal1* ablation in the NAc (n = 7 – 9 per experimental group and sex). Daily alcohol intake (left) and average alcohol intake (right) is increased in *Bmal1-KO* male (A) and female mice (B). Likewise, daily alcohol preference (left) and average alcohol preference (right) is increased in *Bmal1-KO* male (C) and female mice (D). No differences were found in sucrose consumption (E) and preference (F) between *Bmal1-KO* or CTR male and female mice. Values are presented as mean ± standard error of the mean (SEM). # … p ≤ 0.05, ## … p ≤ 0.005, ### … p ≤ 0.0005, two-way repeated measure ANOVA. *** … p ≤ 0.0005, **** … p ≤ 0.00005, unpaired t-test. Details of the statistical tests are depicted in Tables 1 + 2.

**Table 2:**
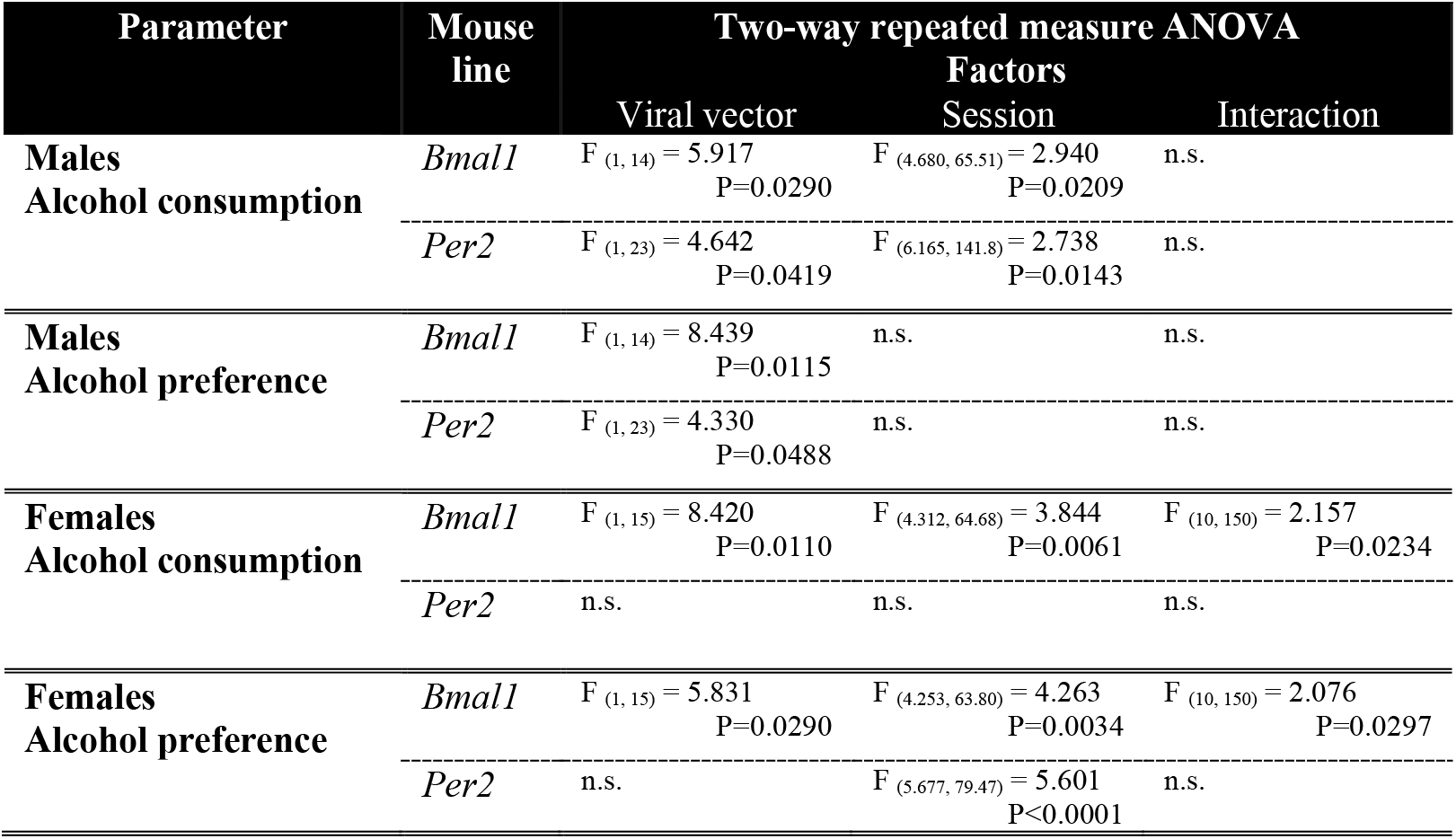
Results of alcohol consumption experiment.

### Effect of *Per2* deletion in the Nac on anxiety and depressive-like behavior

Deletion of *Per2* from the Nac affected anxiety-like behavior displayed in the EPM in a sex-dependant manner. The proportion of time spent in the open arms was affected by the knockout, but the response varied between males and females (Table 1). Whereas a significant increase was found in females, the opposite trend was observed in males (Fig. 4 A). A sex-effect was observed for total distance travelled in the EPM, suggesting that females displayed less activity in the EPM compared to males (Fig. 4 C). Moreover, sex effects were found in behaviors studied in the open field, where males spent more time in the central area of the open field despite a longer latency to enter it (Fig. 4 D). Marble burying was not affected by the *Per2* deletion in either males or females (Fig. 4 G). In the TST, a sex-dependent effect of viral treatment on immobility time was found, indicating that mobility time is decreased in KO females by the Nac-specific deletion of *Per2* (Fig. 4 H, Table 1). Females, however, displayed a strongly reduced mobility time when compared to males in general (Fig. 4 H, Table 1).

**Figure 4:**
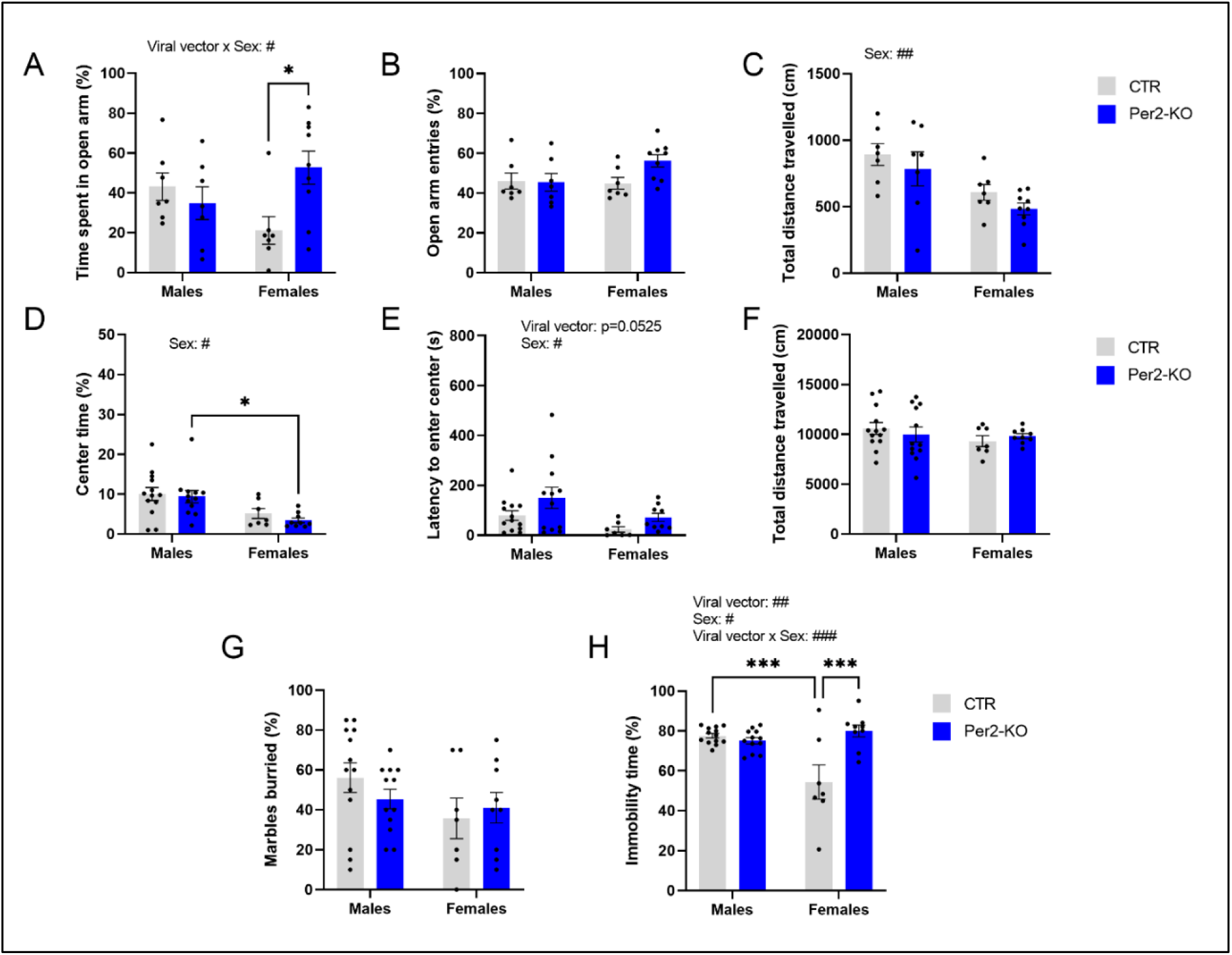
Anxiety- and depressive-like behavior in mice with a NAc-specific *Per2* knockout (n = 7 – 13 per experimental group and sex). While females tended to be less active in the elevated plus maze (C), time spent in the open arm (A) but not open arm entries (B) were significantly increased in *Per2-KO* animals compared to controls. In contrast, center time in the open field was reduced in females irrespective of the administered viral vector (D). The latency to enter the central area of the open field (E) tended to be increased, and males took longer to approach the center compared to females. No differences between groups were found in the marble burying test (G). *Per2*-CTR females were significantly more active in the tails suspension test, which decreased through the NAc-specific deletion and reached comparable levels to males (H). Results are depicted as mean ± standard error of the mean (S.E.M.). # … p ≤ 0.05, ## … p ≤ 0.005, ### … p ≤ 0.0005, two-way ANOVA. *… p ≤ 0.05, **… p ≤ 0.005, Šídák’s multiple comparisons test. Details of the statistical analysis are shown in Table 1.

### Effect of *Per2* deletion in the Nac on alcohol and sucrose consumption and preference

Deletion of *Per2* in the Nac augmented alcohol intake and preference in males, but only gradually and to a lesser extent than *Bmal1* deletion, whereas ethanol consumption or preference was not affected in females (Fig. 5 A – D, Table 2). On average, Nac-KO males drank 3 g/kg more ethanol and displayed 12% higher ethanol preference than CTR animals (Fig. 5 A + C, right panel). Moreover, the deletion of *Per2* in the Nac had no effect on body weight and on sucrose intake and preference in male and female mice (Fig. 5 E + F), whereas a sex effect on sucrose intake was noted (Fig. 5 E, Table 1).

**Figure 5:**
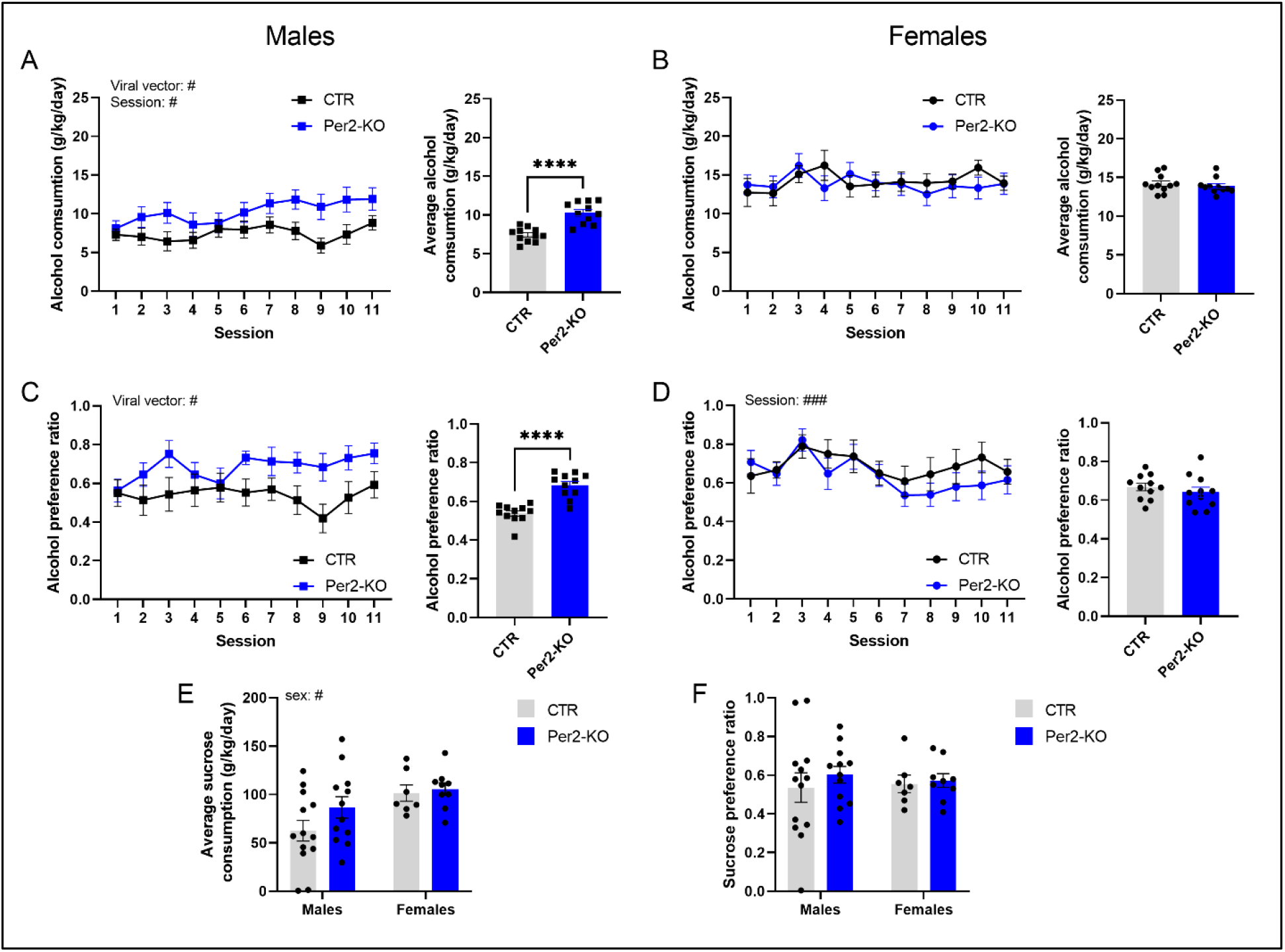
Alcohol and sucrose consumption and preference in mice with a *Per2* deletion in the NAc (n = 7 – 13 per experimental group and sex). Daily alcohol intake (left) and average alcohol intake (right) is increased in *Per2-KO* male (A) but not female mice (B) when compared to CTR animals. Similarly, daily alcohol preference (left) and average alcohol preference (right) is increased in *Per2-KO* males when compared to controls (C), but no difference was observed in females (D). Although females tend to consume more sucrose solution (E), no differences were found between *Per2*-KO or CTR animals. Likewise, sucrose preference was similar between knockout and control male and female mice. Values are presented as mean ± standard error of the mean (SEM). # … p ≤ 0.05, ### … p ≤ 0.0005, two-way repeated measure ANOVA. **** … p ≤ 0.00005, unpaired t-test. Details of the statistical tests are depicted in Tables 1 + 2.

## Discussion

In the present study we examined the Nac as a critical site of sexually dimorphic effects of *Bmal1* and *Per2* on alcohol consumption. AAV-Cre mediated deletion of *Bmal1* in cells of the Nac enhanced alcohol consumption in both male and female mice, whereas deletion of *Per2* in the Nac augmented alcohol intake in males but had no effect on female consumption.

These outcomes largely reflect the results of a previous study where either *Bmal1* or *Per2* were conditionally ablated in MSNs throughout the entire striatum of male and female mice^9^, with one exception. In female mice, *Bmal1* deletion in MSNs within the dorsal and ventral striatum inhibited ethanol consumption, whereas the opposite was found when *Bmal1* was deleted in the Nac exclusively. The results indicate that *Bmal1* and *Per2* regulate voluntary ethanol consumption distinctively within striatal subregions of male and female mice, either through inhibitory or stimulatory functions.

The underlying sex-independent mechanisms of increased ethanol consumption observed in mice with a Nac-specific *Bmal1* deletion are largely unexplored, but multifaceted consequences of such deletion, including altered cell signaling pathways and neurotransmission could be considered. Circadian clock genes have been shown to affect dopaminergic^18,19^, glutamatergic^6^ and GABAergic^20,21^ neurotransmission. Vice versa, acute alcohol exposure affects dopaminergic and glutamatergic signaling in the Nac^22,23^, which may affect clock gene expression^24,25^ and downstream responses through this interaction. Although it cannot be ruled out that the consequences of the clock gene deletion in the Nac and ethanol consumption mutually affect each other, some factors such as sex or the specific clock gene function need to be considered separately. Because Nac-specific *Bmal1* and *Per2* deletion increase voluntary ethanol consumption in males, and because *Bmal1* directly affects *Per2* expression, a shared Nac-specific mechanism could be suggested in males^6^. On the other hand, gene expression studies in the dorsal striatum of conditional *Bmal1* KO mice revealed only minor reductions in *Per2* mRNA levels in KO mice^16^, raising the possibility that both genes act independently of each other. In females, however, this mechanism appears to be strictly *Bmal1*-dependent as Nac- *Per2* KO females do not show changes in voluntary ethanol consumption.

Female sex hormone signaling could further contribute to the observed ethanol drinking phenotypes. The MSNs in the striatum express membrane-associated estrogen receptors, and estradiol has been shown to affect striatal MSN signaling capabilities in a sex- and region-specific manner^26,27^. In addition, estrogen receptor beta (Erβ) gene expression is controlled through BMAL1-CLOCK^28^ which may explain differences in ethanol consumption in female mice with a Nac-specific *Bmal1* or *Per2* deletion. Although Erβ appears to be comparatively little expressed in the Nac, it was found to selectively control responses to drugs of abuse^29^. However, a link between Erβ expression and alcohol consumption has yet not been established and a detailed analysis of the sex-hormone signaling pathways is required to support this hypothesis.

Our previous and present studies of the role of striatal *Bmal1* on alcohol consumption point to an important interaction between sex and striatal subregion. In females, deletion of *Bmal1* in MSNs within the entire striatum has been shown to inhibit ethanol intake, whereas in the present study, *Bmal1* deletion in the Nac promoted consumption. In contrast, in males, deletion of *Bmal1* in the entire striatum or in cells of the Nac only result in the same drinking pattern. Together, these findings mark the Nac as site of sexually unbiased action of *Bmal1* on alcohol consumption and further emphasize a critical role of the dorsal striatum as site of sex-dependent influence of *Bmal1* on alcohol consumption.

While it is logical to assume that the differences in observed ethanol-drinking phenotypes in females with a *Bmal1* knockout in the entire striatum and *Bmal1* ablation restricted to the Nac originate from a region-specific effect of the gene, it is also important to consider other factors, such as the age-dependent effects of the gene knockout itself. Deletion of *Bmal1* in the entire striatum through crossbreeding with the Cre driver line may disrupt development and neurogenesis of MSNs in the Nac and cause long-term alterations which are different to the effects of an ablation through Cre-expressing viral vectors in adult mice. *Bmal1* has been shown to affect early neurogenesis in the brain^30^. Also, it has been suggested that estrogen signaling through both nuclear receptor subtypes affects striatal differentiation^31^, and given the interaction of BMAL1-CLOCK with Erβ gene expression, it may explain differences obtained in females with a region-specific or full striatal *Bmal1* knockout. On the other hand, it has been indicated that the striatal clock is only fully functional in adult individuals, and that levels of *Bmal1* and *Clock* mRNA expression gradually increases, with the lowest levels at postnatal day 3 in mice^32^. Therefore, age-dependent effects of *Bmal1* on MSNs function should be studied systematically at various ages to get a conclusive picture.

Besides the role of striatal MSNs, other cell types in the striatum have been shown to mediate effects of alcohol. Although they represent only 3-5% of the striatal neuronal population, fast spiking interneurons appear to modulate striatal output in response to ethanol^33,34^. Glial cells such as astrocytes, on the other hand, appear not to play a central role in the observed phenotypes even though they have been identified to affect certain aspects of alcohol-drinking behavior^35^. The specificity of the conditional clock gene knockout through crossbreeding or viral approaches, however, make their contribution to the observed phenotypes unlikely. Nevertheless, further region- and cell-type specific approaches are required to identify the exact location and neuronal populations controlling ethanol drinking in striatal clock-KO mice.

Alcohol consumption is influenced by emotional states^36^, and emerging evidence suggest an important role of clock genes in emotion regulation^37^. To identify potential links between emotional states and alcohol drinking, affective behaviors were assessed in mice of the current study before they were exposed to alcohol. Anxiety and depressive-like behaviors were only mildly affected, and sucrose consumption was not altered by Nac deletion of *Bmal1* or *Per2* in either males or females, suggesting that the differences in alcohol intake may involve specific clock and sex-specific mechanisms associated with alcohol-drinking behavior rather than alterations in affective or appetitive states of the animal. This is in accordance with results observed in male and female mice with a *Bmal1* or *Per2* deletion in the entire striatum, which revealed only mild changes in affective behaviors and no difference in sucrose preference in either males or females^9,16^.

Taken together, we hypothesise that the presence of *Bmal1* and *Per2* in the striatum inhibit voluntary ethanol consumption in males, presumably through a common, Nac-specific mechanism. In females, only *Bmal1* appears to have the same inhibitory function in the Nac as demonstrated by the results of the Nac-specific ablation in this study. Strikingly, the effect is inverted when *Bmal1* is conditionally knocked out from MSNs throughout the entire striatum. It suggests a female-specific stimulatory effect of *Bmal1* in the dorsal striatum on voluntary alcohol consumption, which prevails over the inhibitory function in the Nac.

## AUTHOR CONTRIBUTIONS

SA and KS conceived and designed the study. JH performed the experiments with assistance from MB, PD-H, CG, and NQ. JH, SA and KS analyzed the data and prepared graphs. SA and KS wrote the manuscript. JH revised the manuscript. All authors approved the final manuscript.

